# Advancing the Application of pXRF for Biological Samples

**DOI:** 10.1101/2024.01.16.575873

**Authors:** K.J. Brandis, R. Francis, K.J.A. Zawada, C.D. Hasselerharm, D. Ramp

## Abstract

Point 1: Portable x-ray fluorescent (pXRF) technology provides significant opportunities for rapid, non-destructive data collection in a range of fields of study. However, there are sources of variation and sample assumptions that may influence the data obtained, particularly in biological samples.

Point 2: We used representative species for four taxa (fish, mammals, birds, reptiles) to test the precision of replicate scans, and the impact of sample thickness, sample state, scan location and scan time on data obtained from a pXRF.

Point 3: We detected significant differences in concentration data due to sample state, scanning time and scanning location for all taxa. Infinite thickness assumptions were met for fish, reptile and mammal representatives at all body locations when samples were thawed, but not dried. Infinite thickness was not met for feathers. Scan time results found in most cases the 40, 60 and 80 second beam times were equivalent. Concentration data across replicate scans were highly correlated.

Point 4: The opportunities for the use of pXRF in biological studies are wide-ranging. These findings highlight the considerations required when scanning biological samples to ensure the required data are suitably collected, while maintaining minimal radiation exposure to live animals.

## Introduction

Elemental and isotopic signature analyses are routine in the biological and environmental sciences, being used as indicators of diet (Hobson 1992, Pearson et al. 2013), ecotoxicology (Gashkina 2017), pollution (Marcussen et al. 2008), soil sciences (Mancini et al. 2019) and animal movement studies (Hobson 1999, Brandis et al. 2021). Current analytical techniques include stable isotope analysis (SIA), inductively coupled plasma mass spectrometry (ICP-S), atomic absorption spectroscopy (AAS), and neutron activation (NAA) among others (Helaluddin et al. 2016). However, many of these techniques are destructive, time consuming, and expensive, precluding their use in the field, or on live organisms. Development of rapid, *in situ*, inexpensive and non-destructive sampling techniques would greatly expand the utility of elemental signatures in biological and environmental sciences.

X-ray fluorescence (XRF) is a well-established methodology for measuring the elemental composition of samples (Beckhoff et al. 2007). It is a quantitative, non-destructive technique that measures the abundance of an element based on the characteristic emission of secondary x-rays following excitation by a primary x-ray beam (Gates 2006). Portable x-ray fluorescence (pXRF) instruments provide opportunities for field-based data collection. Developed initially for use in the field of geology (Lemière 2018), they are also used in archaeology (Newlander et al. 2015, Mauran et al. 2019) and art history (Lopez 2017, Molari and Appoloni 2021) among others (Hou et al. 2004). Data obtained from pXRF is also used for a range of applied biological questions including the detection of disease (Estevam and Appoloni 2013), toxicology (McInver et al. 2015), and geographic provenance (Buddhachat et al. 2017). However, it’s use in non-human biological studies has been limited.

The portability, non-destructive and rapid multi-variable data collection offers significant opportunities in the field of biological sciences. However, due to its initial development for use in geology there are assumptions about sample characteristics that are not necessarily met by biological, and other non-geological samples. This has potential implications for the way in which raw data is processed by pXRF on-board algorithms and the data provided to the user.

These assumptions include the thickness and density of the sample, referred to as ‘infinite thickness’, i.e., the sample is of sufficient thickness and density that all x-rays are contained within the sample and do not pass through it (Kieser and Mulligan 1979, Sitko 2009). It has been recognized that data acquired from samples that are thin, and do not meet infinite thickness need to be corrected (Sitko 2009). The thickness required to meet this assumption varies with the density of the material being analysed (Liu et al. 2018), which is challenging to accurately determine in living samples.

Moisture content is another variable that can impact on the results returned by pXRF (de Santana et al. 2019, Padilla et al. 2019). Water absorbs the characteristic x-rays from the elements and causes the primary radiation from the excitation sources to scatter. This results in a decrease in the intensity of characteristic x-rays and an increase in the intensity of scattered x-rays in a fluorescence spectrum (Ge et al. 2005). Higher moisture content in soils has been shown to impact data resulting in lower elemental concentrations (Padilla et al. 2019). Moisture content of biological samples is highly varied and dependant on sample state, and in living samples moisture content is very difficult to determine.

The length of time a sample is exposed to x-rays can impact on the data obtained. Scanning times may affect the ability to detect low-concentration or low atomic weight elements, where longer scan times improve the signal-to-noise ratio (Zhang et al. 2021). However, in a biological context, there may be a trade-off between detecting elements, and exposing a live organism to excessive radiation. As such, determining whether elemental signatures significantly change as a function of scanning time is a key consideration to optimise data quality and organism welfare.

Further, the location of the scan on the sample can also influence results. The partitioning of elements in biological samples may occur due to tissue type, metabolic processes (Tomlinson et al. 2004), age, and sex (Underwood 1977). Studies by Nganvongpanit et al. (2016) found different elemental concentrations for different bones within the same individual while Buddhachat et al., (2016) found differences between tissue types with keratinous tissues (hair, nail, skin), being different to bone.

The use of portable X-ray fluorescence (pXRF) as a non-destructive technique for elemental analysis of biological samples offers significant opportunities in the field of biological and environmental sciences. However, when planning a sampling design for biological samples, there are a number of potential sources of variation that need to be accounted for to ensure data are comparable and research questions are answerable. This paper aimed to explore the application of pXRF for elemental analysis of biological samples and to test the major sources of variation across four representative taxa, namely infinite thickness, sample state (dried/thawed), scanning time, and scanning location on the sample. The findings of this study provide insights for future studies and will contribute to the development of standardised protocols for pXRF analysis of biological samples.

## Methods

### Sampling design

We chose species representing four taxa commonly studied in biological research: reptiles, mammals, birds and fish. Representative species for each taxa were chosen based upon accessibility to sufficient specimens, with each species represented by ten samples.

Reptiles were represented by the shingleback lizard (*Tiliqua rugosa*) and were sourced as whole or gutted carcasses from Taronga Conservation Society, Macquarie University and Murdoch University. Mammals were represented by the European rabbit (*Oryctolagus cuniculus*) and sourced from a pet food supplier. Fish were represented by Eastern school whiting (*Sillago flindersi*) sourced from the Sydney fish markets, Australia. Birds were represented by Australian maned duck (*Chenonetta jubata*), and moulted flight feathers were collected from wetland sites. All taxa except the duck feathers were stored frozen prior to scanning. Each of the taxa were used to test each of the sources of variation.

### pXRF data collection

An Olympus Vanta M-Series portable x-ray fluorescence (pXRF) instrument (4-watt X-ray tube with tungsten (W) anode, 8-50keV, silicon drift detector), with three beam energies (10, 40 and 50keV) was used to scan all specimens. Where size permitted, specimens were scanned within the Olympus workstation (Olympus 2020). Samples were scanned using the GeoChem3 method which uses a fundamental parameters algorithm (Afonin et al. 1992) that automatically corrects for inter-element effects (Olympus 2020). The Vanta pXRF provides two output data types; raw beam spectra containing information across 2048 keV bands, and elemental concentrations as a percentage for 42 elements calculated via Olympus’ on-board algorithms.

Reptile, mammal, and fish specimens were scanned in both thawed and dried states, feathers were scanned in their natural state. All specimens were scanned at 20, 40, 60, and 80s/beam with and without silica backing at multiple locations (Figure 1) and states (Figure 2). The silica dioxide block backing consisted of 30 mm diameter x 25 mm solid cylinder. Prior to scanning, specimens were defrosted at room temperature for ∼12 hours and scanned thawed. Specimens were then oven dried (60^°^C for 72 hours) and re-scanned.

**Figure 1.**
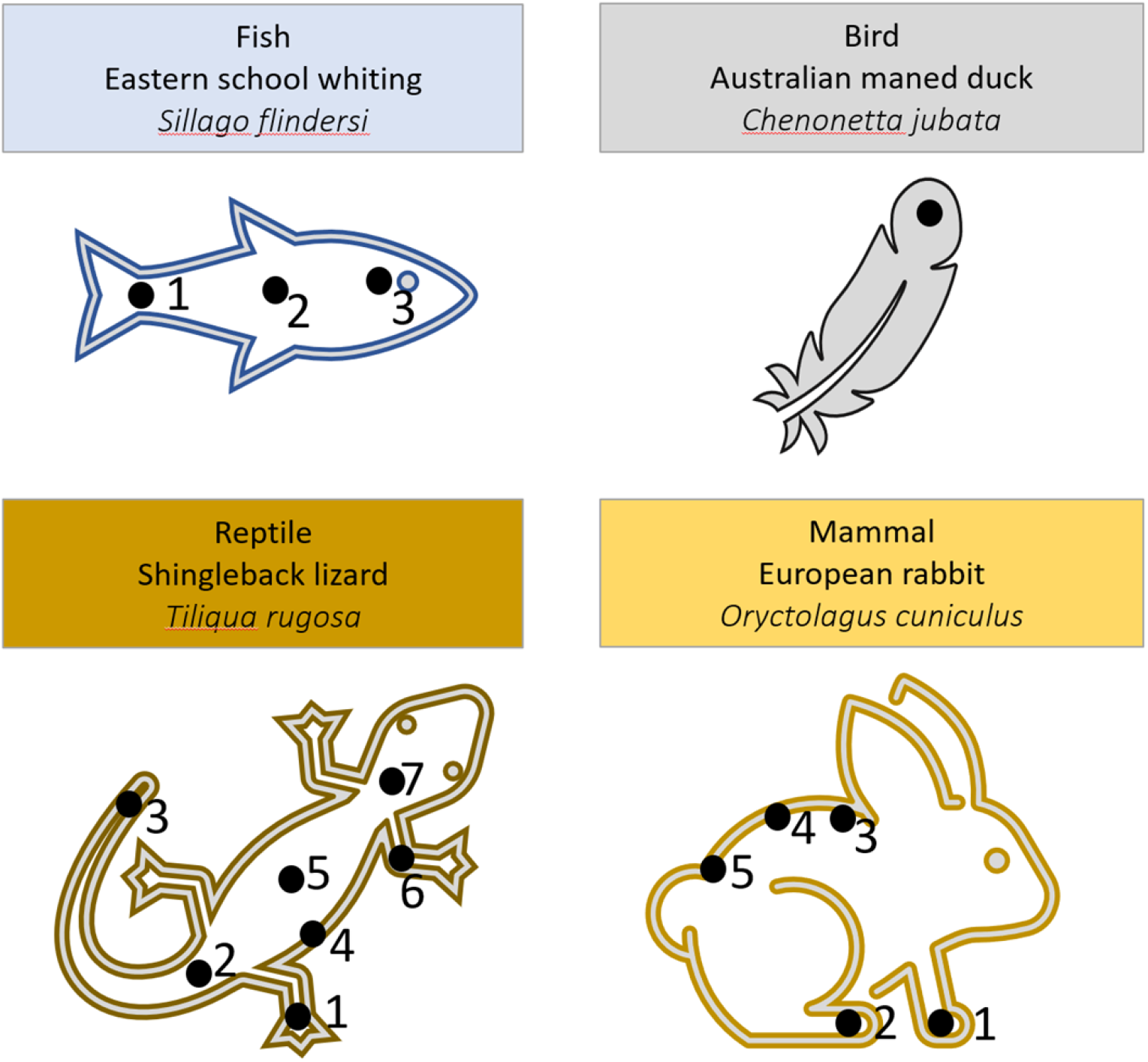
Scanning locations (black dots) on each specimen of representative taxa.

**Figure 2.**
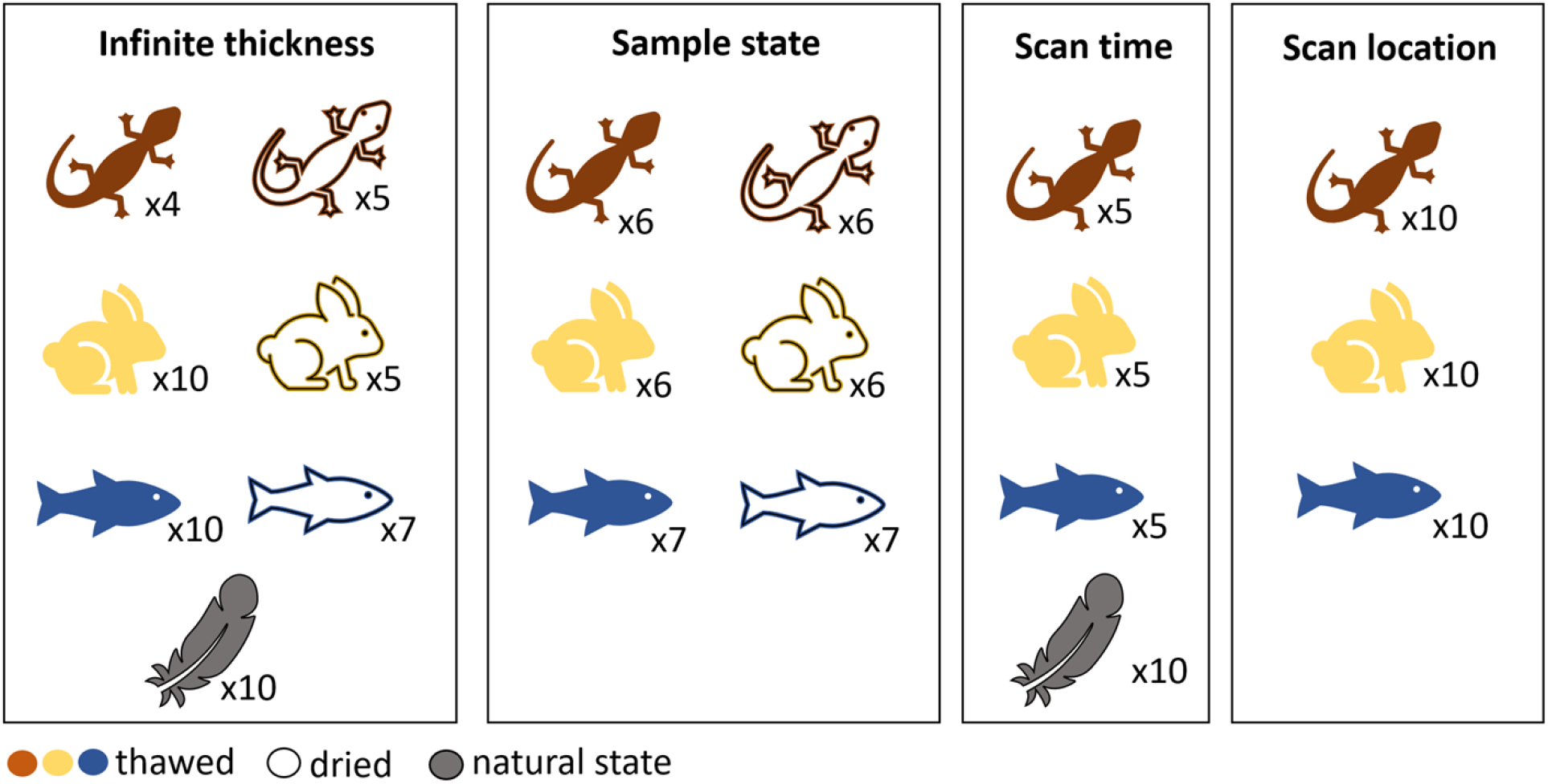
Summary of the number of replicates per variation test and the condition of the specimen; natural state, thawed or dried.

### pXRF data analyses

To test whether infinite thickness was being met we compared the measured silica signal between the silica backed and non-backed samples for 20s/beam scans. If infinite thickness was being met, then no additional silica signal should ‘leak’ into the sample scan when comparing silica and non-silica backed samples. However, if the detected silica concentration increased for the silica-backed samples, this indicated that the outgoing x-rays were passing through the sample and striking the silica block, resulting in a higher detected silica signal. Beam spectra were also analysed as infinite thickness may be met for some beam energies and not others e.g., a 50 keV beam will penetrate further than a 10 keV beam.

We fitted a median quantile regression for each scan location for each species, with silica signal as the response and backing status (silica-backed vs no-backing) as the fixed variable. Median quantile regression was chosen as it is more robust than standard least-squares linear regression to skew and kurtosis in the residuals (Koenker 2005).

Sample state, scanning time, and scan-location were all tested using the same approach using the concentration data (20, 40, 60 and 80s/beam), and were split into an overall difference test across all elements and a per-element analysis to identify specific element differences. In cases with multiple response categories such as scanning location or scanning time, pair-wise models were run to, for example, compare leg to tail, leg to head, head to tail, etc. Per-element analyses followed the median quantile approach described in the infinite thickness section above. Overall differences were tested using multi-response generalised linear models (Wang et al. 2012), or manyglm’s. Manyglm’s allow a model to be specified with multiple response variables (in this case the concentration of each element in a sample) and to calculate an overall effect of an explanatory variable such as tail vs head scanning location. This approach examines all elemental differences simultaneously, and provides overall statistics, including an overall p.value for the effect of the explanatory variable. Unfortunately, no median quantile version of a manyglm is available at present, meaning that these models rely on the assumptions of linear regression. Despite this, they are still likely more robust than typical distance-based approaches (Wang et al. 2012).

### Precision testing

We also tested for correlation between replicate scans to determine the precision of the pXRF when working with biological samples, using Spearman’s correlation coefficient.

## Results

Overall, we detected significant differences in concentration data obtained due to sample state (Figure 3, Supplementary Dataset 1), scanning time (Figure 3, Supplementary Dataset 2) and scanning location for all taxa (Figure 3, Supplementary Dataset 3). All three taxa (reptiles, fish, and mammals) tested for differences between dried and thawed sample states found significant differences in the concentration data overall, with multiple per-element differences identified (Figure 3, Supplementary Dataset 1).

**Figure 3.**
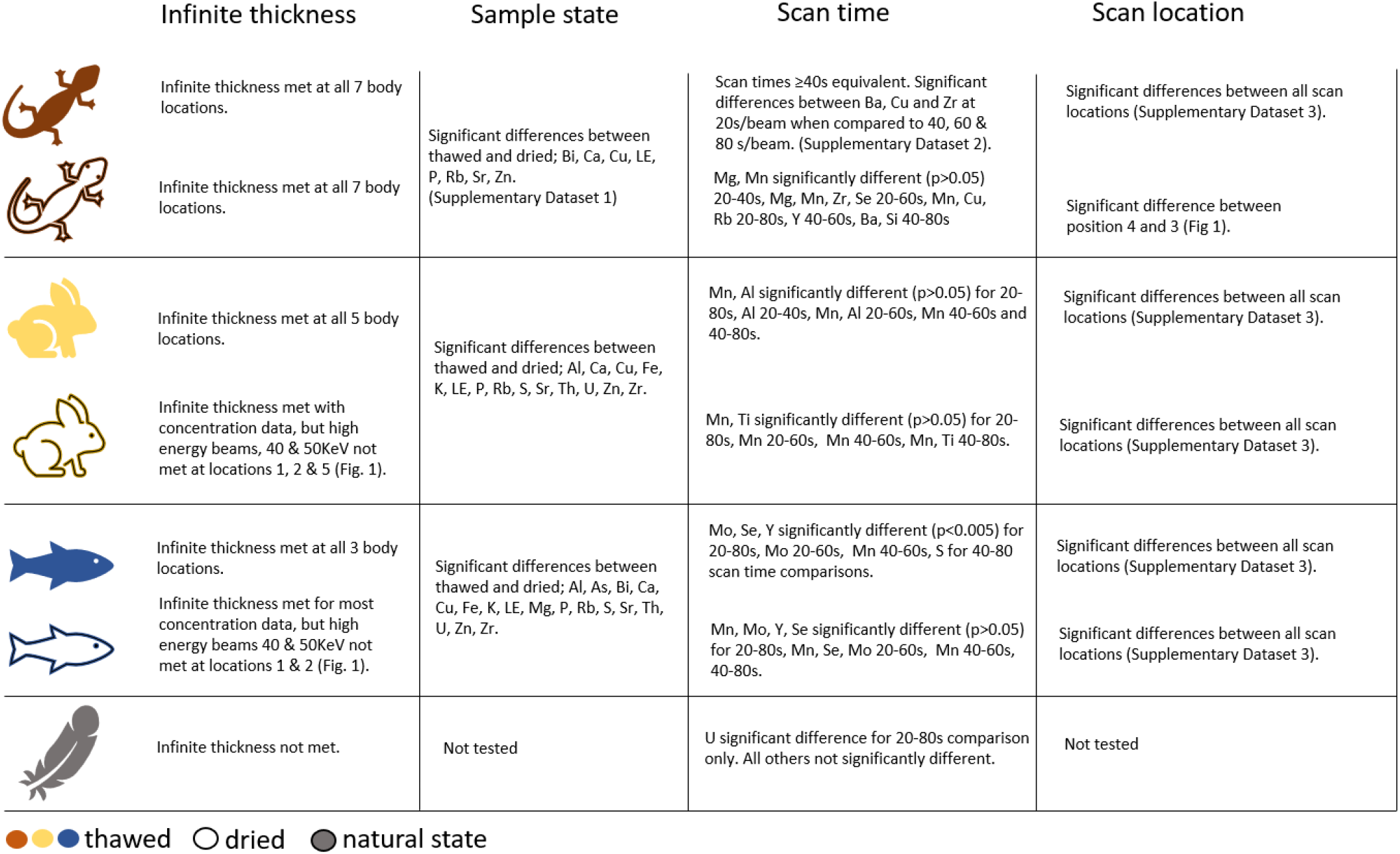
Summary of results for each test using concentration data.

Scanning time had a significant effect on overall detected elemental concentrations for all taxa when comparing the 20 second beam time to the 40, 60 and 80 second beam times, although the differences in the feathers was only for Uranium (Figure 3). All three taxa tested for differences between scanning locations revealed major differences in overall elemental concentration data amongst differing scan locations (Figure 3).

Infinite thickness assumptions were met for thawed states for fish, rabbits and lizards using concentration data, but were not met for some sample states and scanning locations using beam spectra data (Table 1). Infinite thickness was not met for feathers using both concentration and beam spectra data (Table 1).

**Table 1.** Significant results for infinite thickness testing. ^#^cps = counts per second. Asterisks (*) indicate significant differences at the 0.05 (*) and 0.01 (**) level.

| Species | State | Scan location (Fig. 1) | Variable | p.value | Significance |
| --- | --- | --- | --- | --- | --- |
| Rabbit | Thawed | Body base (5) | 50keV.cps#_mean | 0 | ** |
| Rabbit | Thawed | Body base (5) | 40keV.cps_mean | 0 | ** |
| Rabbit | Thawed | Back foot (2) | 50keV.cps_mean | 0 | ** |
| Rabbit | Thawed | Back foot (2) | 40keV.cps_mean | 0.001 | ** |
| Rabbit | Thawed | Front foot (1) | 50keV.cps_mean | 0 | ** |
| Rabbit | Thawed | Front foot (1) | 40keV.cps_mean | 0 | ** |
| Shingleback | Dried | Tail tip (3) | 40keV.cps_mean | 0.025 | * |
| Fish | Dried | Tail (1) | 50keV.cps_mean | 0 | ** |
| Fish | Dried | Body middle (2) | 50keV.cps_mean | 0.016 | * |
| Fish | Dried | Body middle (2) | 40keV.cps_mean | 0.003 | ** |
| Fish | Thawed | Tail (1) | 50keV.cps_mean | 0.049 | * |
| Bird | Natural | Feather tip | 50keV.cps_mean | 0 | ** |
| Bird | Natural | Feather tip | 40keV.cps_mean | 0 | ** |
| Bird | Natural | Feather tip | 10keV.cps_mean | 0 | ** |
| Bird | Natural | Feather tip | concentration | 0.01 | * |

Overall device precision was very high, with a Spearman’s correlation coefficient of 0.999 when comparing replicate scans (Appendix 1). There was variation in the correlation of replicate scans across elements, with heavier elements less correlated than lighter elements (Appendix 2).

## Discussion

In this study we investigated the application of pXRF for sampling biological samples, focusing on the effects of sample thickness, sample state, scanning time, and scan location on elemental concentration data. Our results found that sample state, and scan location resulted in the most significant differences between element concentrations. Infinite thickness assumptions were met for most tests regardless of sample state, excluding feather samples. Scan time results found in most cases the 40, 60 and 80 second beam times were equivalent.

These findings are consistent with previous studies that reported the impacts of sample moisture (Nuchdang et al. 2018, Padilla et al. 2019), and sample thickness (Liu et al. 2018, Padilla et al. 2019) on pXRF data collected from soil samples. Extended scan times were not found to result in significant data gains, which is supported by findings by Williams et al. (2020), however is contrary to findings by Zhang et al. (2021), which found extended (5 minute) scan times reduced variability in results when scanning bone through soft tissue. Extending scan times on live specimens increases radiation exposure which can result in health impacts (Preston et al. 2013). Estevam and Appoloni (2013) calculated that a 50 second scan using an x-ray source of 13 and 17 keV resulted in an exposure of 3 mSV.

Based on the results of this study and considering animal care and ethics we would recommend when scanning live specimens that scan times are kept to a minimum while still ensuring adequate data is obtained, our study suggests that 40 s/beam is sufficient to achieve this for most elements (Figure 3). We would also recommend scanning a part of the body that is of sufficient thickness to meet infinite thickness assumptions, but away from key organs to limit the impact of radiation exposure, and to scan with a standardised and consistent backing behind the specimen whenever possible. For the Olympus Vanta this is the provided silicon dioxide block but any other inert, elementally simple material should suffice. This study found that on thawed specimens (akin to live organisms with regards to body thickness and moisture) the majority of scan locations met infinite thickness requirements (Figure 3). However, noting that different scan locations on the specimen may results in different results (Figure 3). Lastly, noting that we found significant differences in data due the state (thawed or dried) of the sample.

There are some limitations of this study that should be considered when translating our results. Firstly, we only tested a limited number of biological samples from a small set of taxa. Our results may not be generalizable to other types of biological samples but are intended as a guide to the impact different samples, sample characteristics, and sampling techniques may have on results. Secondly, we used a single pXRF instrument with a specific configuration and settings. Different instruments or settings may produce different results (Goodale et al. 2012), however the broad effects demonstrated here should hold as the sources of variation are extrinsic to the instrument. Finally, we did not evaluate the long-term stability or repeatability of pXRF measurements for biological samples (Newlander et al. 2015). Future research should address these issues by testing more diverse biological samples and elements, comparing different pXRF instruments and settings, and assessing the quality control and assurance procedures for pXRF analysis.

This study demonstrates that pXRF can be a useful tool for biological research if sample characteristics and sample design are cognisant of the assumptions of the pXRF device. The advantages of pXRF over conventional analytical techniques include its portability, non-destructiveness, speed, and capability to measure many elements simultaneously. These features make pXRF an attractive option for biological studies that require in situ or large-scale analysis of elemental signatures in various types of samples, such as feathers, hair, bones, plants, soils, etc. However, our study also highlights the considerations required when sampling biological samples using pXRF and the potential impact on results obtained, which include where to scan, how long to scan and how to prepare the sample.

## Acknowledgements

B. Jackson, Murdoch University, WA. P. Meagher, Taronga Conservation Society, D. Harasti, NSW Department of Planning and Environment, The Reptile Lab, Macquarie University.

## Author contributions

KB conceived the idea; All authors contributed to methodology; RF, KZ, and CH undertook sample collection and data capture; KZ, RF and CH undertook data analysis; All authors contributed critically to the draft and gave final approval for publication.

## Data availability

## Appendix 1

**Figure 1.**
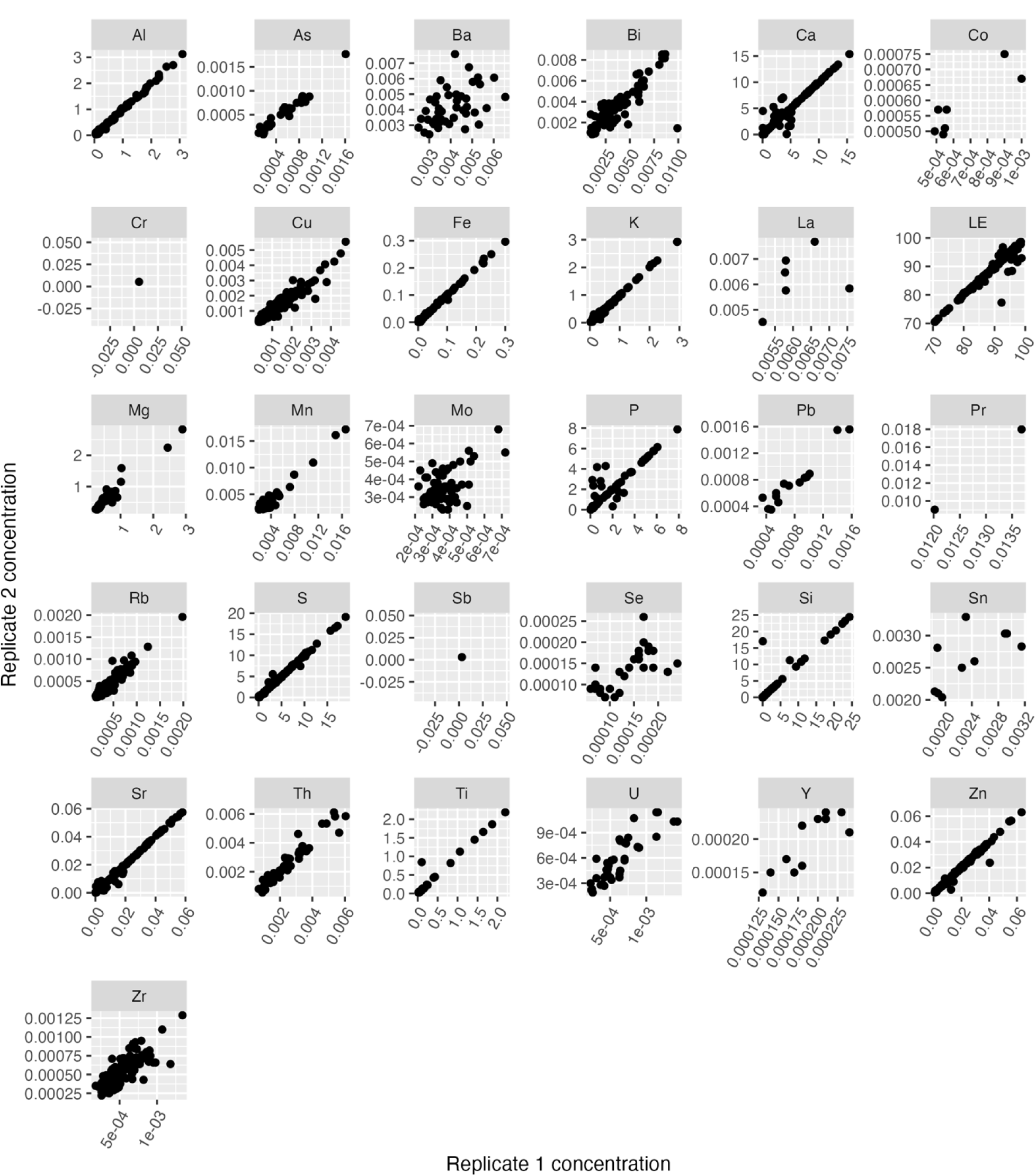
Plot of replicate 1 and replicate 2 scans for each element.

## Appendix 2

**Table 2.** Spearman’s correlation coefficients for each element comparing replicate 1 and replicate 2 scans.

| Element | Spearman's Correlation Coefficient |
| --- | --- |
| La | 0.35 |
| Ba | 0.47 |
| Mo | 0.51 |
| Sn | 0.65 |
| Se | 0.70 |
| Zr | 0.80 |
| Bi | 0.81 |
| Y | 0.85 |
| Co | 0.87 |
| U | 0.87 |
| Si | 0.95 |
| P | 0.95 |
| Mn | 0.95 |
| Cu | 0.95 |
| Rb | 0.95 |
| Mg | 0.96 |
| Pb | 0.96 |
| Th | 0.96 |
| LE | 0.97 |
| Ca | 0.98 |
| Ti | 0.98 |
| As | 0.98 |
| Zn | 0.99 |
| Al | 1.00 |
| S | 1.00 |
| K | 1.00 |
| Fe | 1.00 |
| Sr | 1.00 |

